# Ischemic tolerance and cardiac repair in the African spiny mouse

**DOI:** 10.1101/2021.01.07.425783

**Authors:** Tim Koopmans, Henriette van Beijnum, Elke F. Roovers, Divyanshu Malhotra, Antonio Tomasso, Jochem Boeter, Danielle Versteeg, Eva van Rooij, Kerstin Bartscherer

**Affiliations:** Hubrecht Institute for Developmental Biology and Stem Cell Research, Royal Netherlands Academy of Arts and Sciences, Utrecht, The Netherlands

## Abstract

Ischemic heart disease and by extension myocardial infarction is the primary cause of death worldwide, necessitating regenerative therapies to restore heart function. Current models of heart regeneration are restricted in that they are not of adult mammalian origin, precluding the study of class-specific traits that have emerged throughout evolution, and reducing translatability of research findings to humans. Here, we overcome those restrictions by introducing the African spiny mouse (*Acomys spp*.), a murid rodent that has recently been found to exhibit *bona fide* regeneration of the back skin and ear pinna. We show that spiny mice exhibit tolerance to myocardial infarction through superior survivability, improved ventricular conduction, smaller scar size, and near-absence of cardiac remodeling. Critically, spiny mice display increased vascularization and cardiomyocyte expansion, with an associated improvement in heart function. These findings present new avenues for mammalian heart research by leveraging unique tissue properties of the spiny mouse.

## Introduction

Heart failure affects more than 26 million people worldwide ^1^, making effective therapies for heart failure a major public health goal. A key underlying cause of heart failure is the inability of the adult human myocardium to regenerate following injury. Correspondingly, a central focus in cardiac regenerative research has been the restoration of heart lineages, including vascular cells, support cells, and most notably the adult cardiomyocyte. However, its effectiveness has been limited by the absence of naturally regenerating adult mammalian organisms ^2^, restricting research to either non-mammalian vertebrates (such as zebrafish ^3,4^) or neonatal mice, which are capable of scar-free cardiac repair up to one week after birth ^5^. The African spiny mouse (*Acomys spp*.) has recently emerged as a new model system, primarily due to its ability to scarlessly regenerate full thickness excisional wounds in the back skin and ear pinna ^6,7^. Given the presence of additional anatomical features - i.e. tail sheath autotomy and fragile attachment of the back skin to the underlying fascia ^6^ - the regenerative capacity of *Acomys* likely developed locally in surface tissues as an adaptation to evade predators. However, whether regenerative capacity extends towards the essential internal organs, particularly the heart, is incompletely understood ^8^. Here, we compared the response to ischemic injury following ligation of the left anterior descending (LAD) coronary artery (resulting in myocardial infarction (MI)), in *Mus musculus* and *Acomys cahirinus*, for whom coronary artery anatomy is largely similar ^9^. We report, despite scar formation and an initial drop in heart function, spiny mice exhibit significant tolerance to infarction, demonstrated by superior survivability, retainment of ventricular conduction, smaller scar size, and reduced remodeling. Importantly, *Acomys* hearts display a partial recovery in contractile output, potentially through increased cardiomyocyte and vascular proliferation.

## Results

To capture the dynamics of cardiac ischemia and potential repair following LAD ligation, we analyzed heart function and tissue samples at different time points up to 100 days after initial injury (Fig. 1a). Remarkably, *Acomys* was extremely tolerant towards the surgical procedure and all animals survived, contrary to *Mus* who exhibited 37.5% lethality (Fig. 1b). Yet, *Acomys* developed scar tissue throughout the damaged area, which appeared grossly similar to that of *Mus* (Ext. fig. 1a). We proceeded to accurately quantify scar area using a series of transversal stacks, from point of ligation down to the apex (see ‘Methods’). Whereas *Mus* developed a large scar that continued to expand due to compensatory hypertrophy and progressive dilatation, *Acomys* late stage scars quickly plateaued in relative area, maintaining a smaller size compared to *Mus* (Fig. 1c,d). In accordance with increased scar area, echocardiography revealed a continued worsening of ejection fraction (from 32.1% ± 1.6 at 4 days after MI to 21.5% ± 1.9 at 100 days after MI) and fractional shortening in *Mus*, whereas *Acomys* continuously improved over a 100-day period post-MI (from 39.9% ± 2.2 at 4 days after MI to 47.0% ± 1.7 at 100 days after MI) (Fig. 1e). Importantly, scar area negatively correlated with ejection fraction in *Mus*, whereas *Acomys* showed a much poorer correlation (Fig. 1f). These results suggest scar tissue in *Acomys* differs from conventional mammalian scars, with functional properties that permit cardiac output. Accordingly, we measured electrical conductance of the heart using electrocardiography (ECG), and measured quantitative differences in QRS behavior that serves as a critical indicator of left ventricular conductivity. Most notably, after MI, we found a strong and enduring increase in both Q amplitude and duration for *Mus*, resembling pathological Q waves in human patients ^10^, with no significant alterations for *Acomys* (Fig. 1g and Ext. fig. 1b). To substantiate our findings, we measured functional heart parameters using a reperfusion injury model, whereby we temporary ligated the LAD for 1 hour, after which arterial blood flow is restored. As expected, we observed a much milder, largely reversible phenotype, although both echo and ECG data showed similar differences between *Acomys* and *Mus* as observed for the MI model (Ext. fig. 1c,d). Collectively, our findings demonstrate that *Acomys* exhibits significant tolerance in the context of cardiac ischemia, and shows partial recovery of heart function.

**Figure 1.**
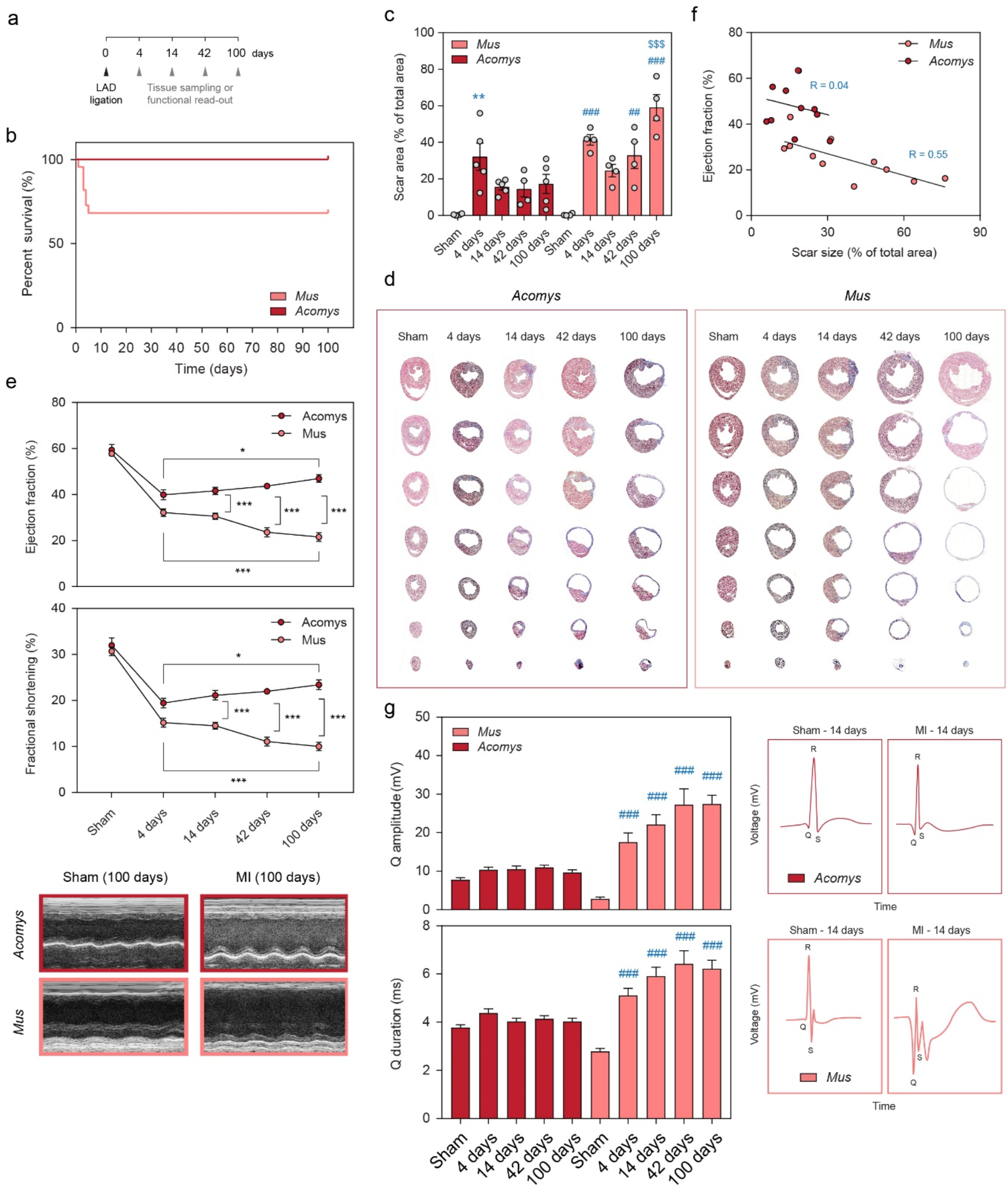
*Acomys* exhibits ischemic tolerance and partially recovers heart function. **(a)** Overview showing the different time points after LAD ligation surgery, used for functional measurements or tissue sampling. **(b)** Kaplan-Meier survival curve for adult *Acomys* and *Mus* after LAD ligation, up to a 100 days post-MI. n = 12 per group. **(c)** Scar area (% of total area) development over time, based on transversal stacks stained with Masson’s Trichrome (see ‘Methods’). * is compared to *Acomys*-Sham, # is compared to *Mus*-Sham, $ represents an inter-species comparison of the same time point (e.g. 100 days to 100 days). One-way ANOVA followed by Tukey’s post hoc test. **(d)** Representative images from (c), showing transversal progression from point of ligation down to the apex. **(e)** Echocardiographic quantification of cardiac output after MI or Sham-control, including ejection fraction (volumetric percentage of fluid ejected from the chamber with each contraction), and fractional shortening (percentage of diastolic dimension that is lost in systole). Corresponding representative images of the M-mode recording are show below the graphs. One-way ANOVA followed by Tukey’s post hoc test. n = 15 for *Acomys*, n = 14 for *Mus*. Graph contains animals that were measured over different time points (paired samples), as well as animals that were only measured at one time point (unpaired samples). **(f)** Correlation plot from all animals that had their ejection fraction and corresponding scar area measured. Graph contains only the animals that had both echo data and scar area measured. **(g)** Electrocardiogram quantification (see ‘Methods’) showing Q wave parameters (amplitude and duration) after MI or Sham-control, with corresponding representative illustration of the QRS wave. Graph contains animals that were measured over different time points (paired samples), as well as animals that were only measured at one time point (unpaired samples). # is compared to *Mus*-Sham. One-way ANOVA followed by Tukey’s post hoc test. n = 15 for *Acomys*, n = 14 for *Mus*. * is p < 0.05, ** is p < 0.01, and *** is p < 0.001.

We next assessed differences in heart and body weight as a proxy for cardiac remodeling (clinically measurable changes that affect heart size, shape or function ^11^). Although *Mus* individuals had a lower total body weight (24 g ± 0.3 compared to 32 g ± 0.6), we observed similar weights for healthy hearts (122 mg ± 2.8 compared to 130 mg ± 4.3) (Fig. 2a). Over time, *Mus* animals took on body weight more rapidly than *Acomys*, but the MI procedure had no add-on effect on body weight compared to Sham-controls, for both species (Fig. 2b). We thus used the body/heart weight metric as a first indication of cardiac remodeling, and observed a stark and continued increase for *Mus*, with only a mild, but reversible increase for *Acomys* (Fig. 2c). Correspondingly, heart size (calculated as the mean surface area from six equally spaced transversal stacks) increased for *Mus*, but not for *Acomys* (Fig. 2d). In addition, mRNA levels derived from whole ventricles marking known targets for heart failure or cardiac hypertrophy ^12^, including members of the myosin heavy chain and natriuretic peptide family, confirmed these findings, showing a dramatic MI-induced increase in *Mus*, but virtual absence in *Acomys* (Fig. 2e). We conclude that *Acomys* hearts are largely protected from pathological remodeling after infarction.

**Figure 2.**
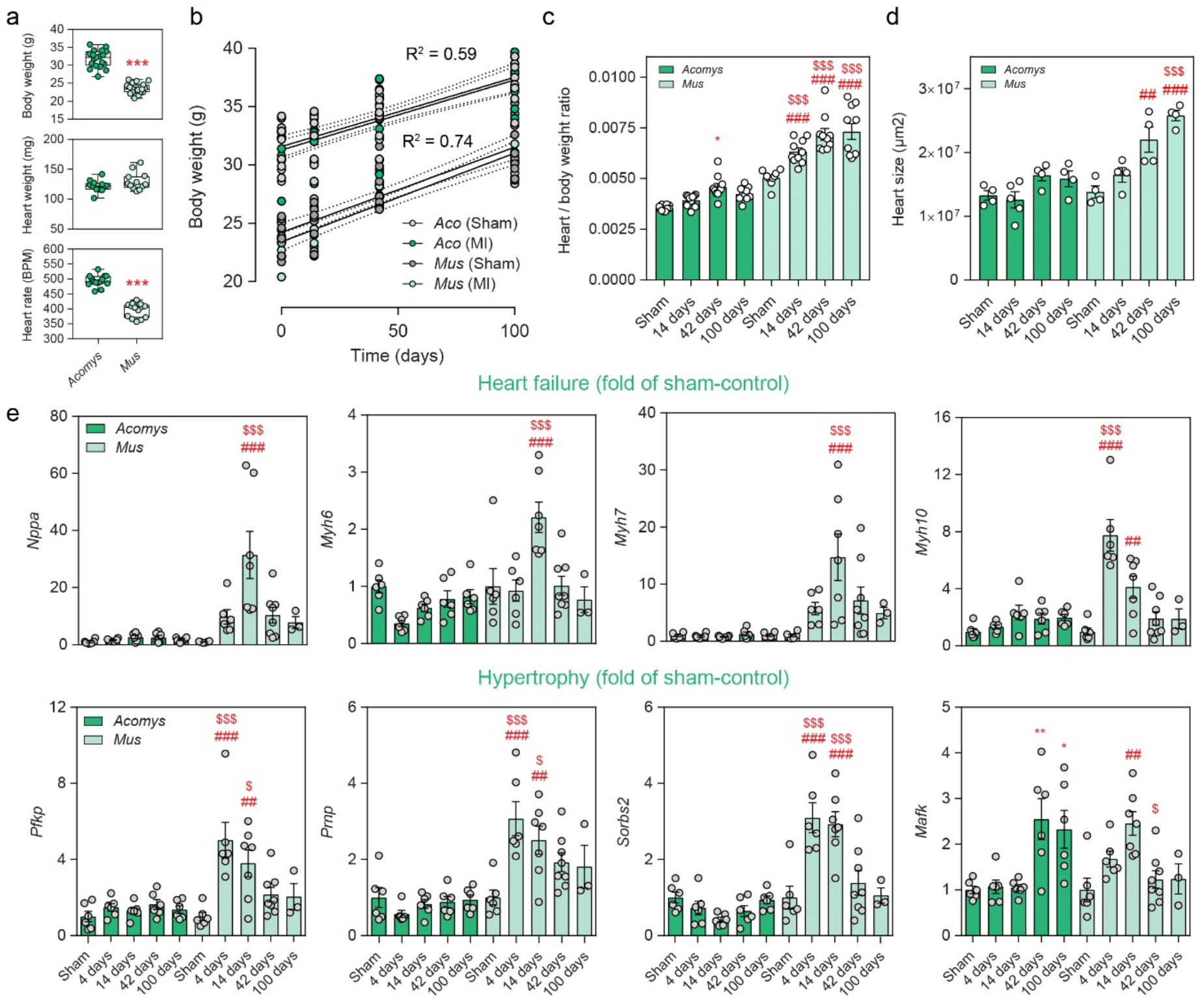
Near-absence of infarction-induced remodeling in *Acomys*. **(a)** Basic metrics comparison between Acomys and Mus. Unpaired t-test. **(b)** Body weight progression after Sham- or MI-surgery. Solid line represents a fitted linear regression. Dashed lines mark the 95% confidence interval boundaries of the best-fit line. **(c)** Heart to body weight ratio progression over time. * is compared to *Acomys*-Sham, # is compared to *Mus*-Sham, $ represents an inter-species comparison of the same time point (e.g. 100 days to 100 days). One-way ANOVA followed by Tukey’s post hoc test. **(d)** Heart size progression, based on transversal stacks stained with Masson’s Trichrome (see ‘Methods’). # is compared to *Mus*-Sham, $ represents an inter-species comparison of the same time point (e.g. 100 days to 100 days). One-way ANOVA followed by Tukey’s post hoc test. **(e)** Relative gene expression from whole heart ventricle homogenates subjected to RT-qPCR. Samples have been normalized to the Sham-control (*Acomys* or *Mus*). * is compared to *Acomys*-Sham, # is compared to *Mus*-Sham, $ represents an inter-species comparison of the same time point (e.g. 100 days to 100 days). One-way ANOVA followed by Tukey’s post hoc test. * is p < 0.05, ** is p < 0.01, and *** is p < 0.001.

To assess a putative capability for cardiac regeneration in spiny mice we first addressed global proliferation dynamics using 5-ethynyl-2’-deoxyuridine (EdU) (Ext. fig. 2a) and subsequent confocal immunofluorescence (IF) imaging. Interestingly, *Acomys* and *Mus* hearts showed similar proliferation dynamics over time within the scar area (Ext. fig. 2b). As proliferation *per se* is not necessarily a measure of cardiac repair, we expanded our search towards specific lineages of the heart.

Accordingly, we first addressed vascular dynamics and angiogenesis. In contrast to *Mus*, we found a large number of expanding EdU+ capillaries in the scar region of *Acomys* MI hearts, marked by isolectin B4 labeling, which remained high up until at least 100 days post-MI (Fig. 3a). In parallel, tissue biopsies taken from either the scar area or healthy myocardium were stained for alpha-smooth muscle actin (α-SMA) to visualize muscularized arteries through whole-mount IF confocal imaging. We observed a large network of vessels in 100 day old *Acomys* scars that appeared well-branched and connected. Conversely, *Mus* scars contained smaller vessels that appeared more fragmented (Fig. 3b). To explore the angiogenic program underlying these responses, we first performed quantitative real time PCR on essential angiogenic factors ^13^. While we did not find significant differences between the species for *Angpt2, Angpt3 and Fgf2, Vegfc* was strongly increased for at least 6 weeks after MI in *Acomys* hearts (Fig. 3c). To reveal additional angiogenic regulators we screened for the expression of 53 different angiogenic proteins in whole ventricles using proteomic profiling. We observed increased expression of a variety of proteins in *Acomys* hearts following injury, including VEGF (Fig. 3d and Ext. fig. 2c). Specifically, IGFBP-2, MMP-8 and MMP-9 showed increased expression in *Acomys* hearts, which we were able to confirm using RT-qPCR (Fig. 3d, e). In summary, although *Mus* initiates an angiogenic program leading to the formation of capillaries and to some extent arteries, the formation of these vessels fails to be maintained over time, or never reaches full maturity to begin with, and is incapable of reconstructing the pre-injury state. By contrast, *Acomys* appears to initiate and maintain a full and mature angiogenic program up to at least a 100 days post-MI.

**Figure 3.**
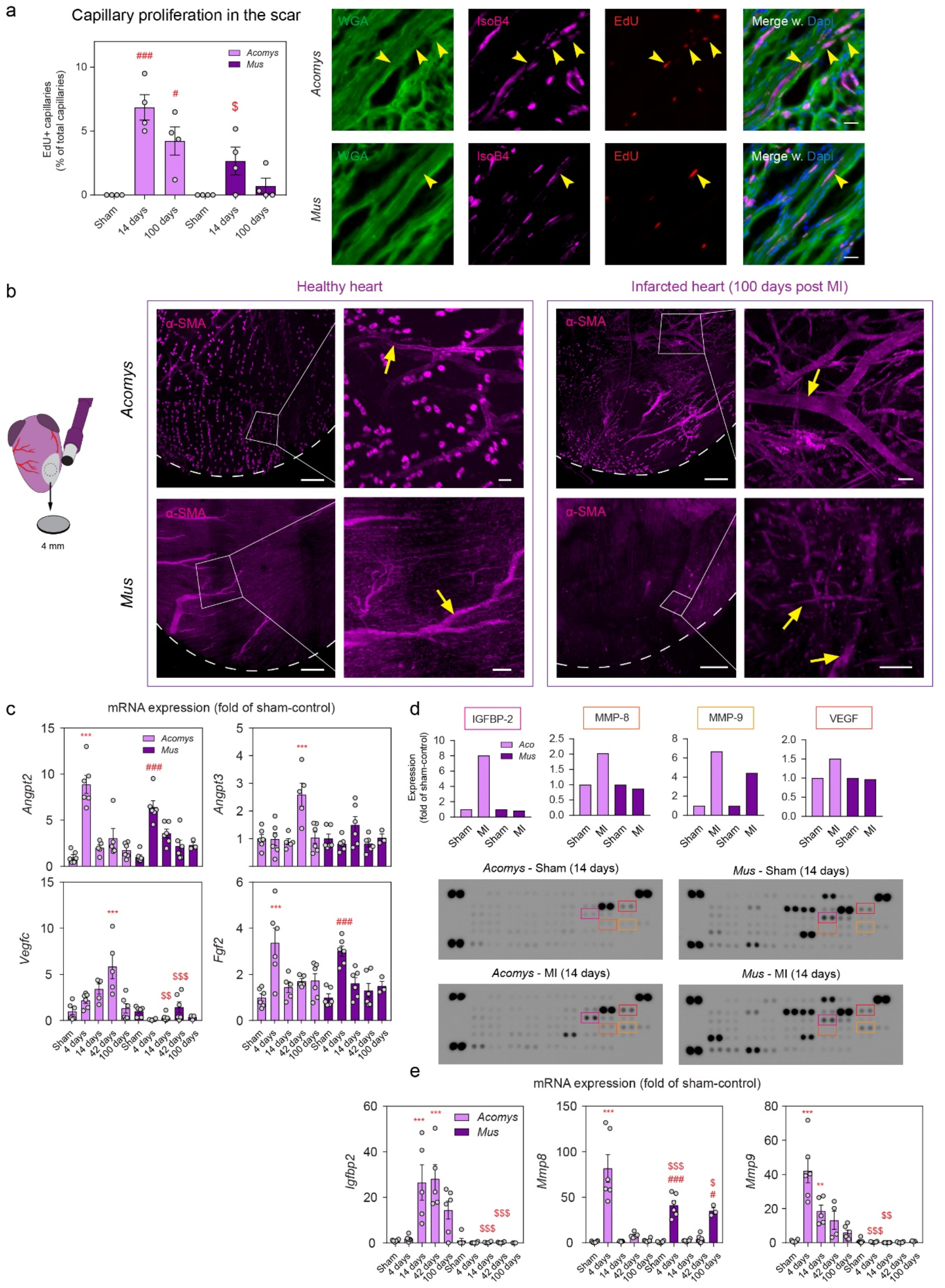
Vascular repair in injured *Acomys* hearts. **(a)** Capillary (marked by Isolectin B4) proliferation in the scar area of infarcted hearts, indicated by EdU labeling. * is compared to *Acomys*- Sham, # is compared to *Mus*-Sham, $ represents an inter-species comparison of the same time point (e.g. 100 days to 100 days). One-way ANOVA followed by Tukey’s post hoc test. Panels at the right side indicate representative fluorescent dye images showing proliferating capillaries (yellow arrows) inside the scar area. Scale bar = 20 µm. **(b)** Whole mount confocal images showing α-SMA+ vessels inside the healthy heart or scar area of 100 day old infarcted hearts. Yellow arrows point towards vessels. Scale bar = 300 µm (overview) or 50 µm (inlet). **(c)** Relative gene expression from whole heart ventricle homogenates subjected to RT-qPCR. Samples have been normalized to the Sham-control (*Acomys* or *Mus*). * is compared to *Acomys*-Sham, # is compared to *Mus*-Sham, $ represents an inter-species comparison of the same time point (e.g. 100 days to 100 days). One-way ANOVA followed by Tukey’s post hoc test. **(d)** Proteomic profiling of angiogenic proteins, from whole heart ventricle homogenates (14 day post-MI and 14 day Sham-controls). Samples are normalized to the Sham-control (*Acomys* or *Mus*). Graphs are derived from the blots pictured below, showing matched colored squares for each indicated graph. Every dot inside the square represents one protein in duplicate. **(e)** Relative gene expression, as in (c). * is p < 0.05, ** is p < 0.01, and *** is p < 0.001.

While improved vascularization by itself may be sufficient to transmit increased cardiac output in *Acomys*, we diverted our attention towards the cardiomyocyte (CM) as the central player of the heart. We first assessed nuclei numbers in isolated CMs as an indication of regeneration potential ^14^, and included zebrafish (*Danio rerio*) in our analysis to establish a ‘regeneration framework’. While *Danio* hearts presented with a single nucleus per cell (1.18 nuclei ± 0.08), and *Mus* with two or more (2.5 nuclei ± 0.10), *Acomys* CMs (2.1 nuclei ± 0.06) combined with their surface area presented as an intermediary in between that of *Mus* and *Danio*, suggesting CMs in *Acomys* exhibit partial immaturity and accompanying proliferative potential ^15^ (Fig. 4a). To further test this hypothesis, we performed bulk sequencing on whole ventricles on adult (7 wks) and neonatal (p3) mice - which still have the potential to regenerate damaged heart tissue ^5^ - to establish maturity parameters of the healthy myocardium, compared between the species. We employed an additive model to compare the effects of age, while controlling for species, and found many differentially expressed genes (Fig. 4b).

**Figure 4.**
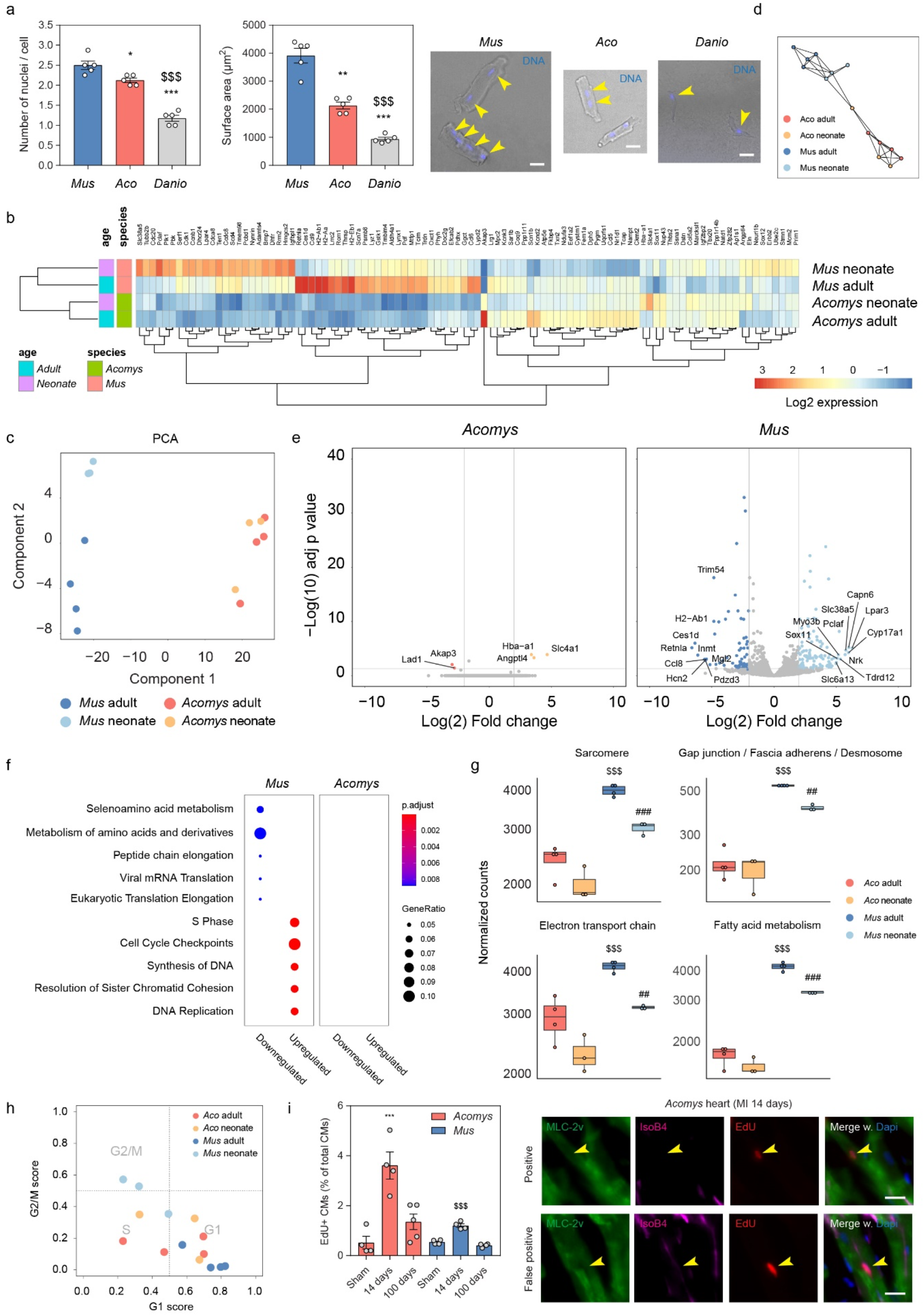
Cardiomyocytes in *Acomys* hearts have an immature phenotype and display increased proliferation post-MI. **(a)** Nuclei number and corresponding surface area of isolated cardiomyocytes (CM) using potassium hydroxide (see ‘Methods’). Panels at the right side indicate representative images of isolated CMs. Yellow arrows point towards multiple nuclei per cell. Scale bar = 30 µm. **(b)** Heatmap from healthy bulk-sequenced heart ventricles, showing the top 100 most significant differentially expressed genes between the samples. **(c)** Sample-to-sample distance visualization through principal component analysis (PCA). **(d)** Pairwise correlation network reveals a maturation topology. Correlation threshold, Pearson’s r > 0.4. **(e)** Volcano plot of top regulated genes (after thresholding) in *Acomys* (adult vs neonate) and *Mus* (adult vs neonate). **(f)** Over- or underrepresented enrichment terms (Reactome database) in *Acomys* (neonate) and *Mus* (neonate). **(g)** Cumulative expression score of genes that fall under different ‘Cellular components’ GO terms, e.g. the sarcomere (GO:0030017). **(h)** Assignment of cell cycle stage for each sample (see ‘Methods’). **(i)** CM (marked by MLC-2v) proliferation that are Isolectin B4 negative (to exclude capillaries) in heart ventricles, indicated by EdU labeling. * is compared to *Acomys*-Sham, $ represents an inter-species comparison of the same time point (e.g. 100 days to 100 days). One-way ANOVA followed by Tukey’s post hoc test. Panels at the right side indicate representative fluorescent antibody or dye images showing proliferating (positive and false positive) CMs (yellow arrows). Scale bar = 15 µm. * is p < 0.05, ** is p < 0.01, and *** is p < 0.001.

Interestingly, principal component analysis (PCA) showed clearly separated clusters for the *Mus* populations, whereas *Acomys* adult and neonate overlapped (Fig. 4c). Using genes from the PCA, we inferred lineage relationships among the samples in an adjacency network on the basis of pairwise correlations between samples ^16^, which confirmed the species separation, with overlap within the *Acomys* pool (Fig. 4d). Correspondingly, when adult hearts were individually compared to neonatal hearts, virtually no significant genes above a 2.0 log_(2)_ fold change threshold were found for *Acomys*, whereas these differences were much more apparent in *Mus* (Fig. 4e). Equally, while we found enrichment terms associated with cell cycling and DNA synthesis for *Mus* neonates, no significant enrichment was found in *Acomys* (Fig. 4f). These results strengthen the hypothesis that adult *Acomys* ventricles exist in a more immature state as compared to *Mus*. To expand on these findings, we generated a cumulative score of genes associated with CM maturation, based on published GO terms (cellular components): 1) sarcomeric load (GO:0030017), 2) cellular integration (GO:0005921, GO:0005916, GO:0030057), 3) electron transport chain components (GO:0022900), and 4) and fatty acid metabolism (GO:0006631) ^17^. Consistently, we found elevated expression in *Mus* adults compared to *Mus* neonates, but an even lower expression for *Acomys* regardless of age (Fig. 4g), further supporting a juvenile phenotype of the ventricular myocardium in *Acomys*. Finally, to establish the potential proliferative capacity of CMs, we predicted cell cycle states based on expression patterns of pre-trained classifiers (see ‘Methods’) ^18^. All *Mus* adult replicates existed in G1 phase, while *Mus* neonates mostly occupied G2/M. Interestingly, *Acomys* adult and neonates displayed a shift from G1 into S phase (Fig. 4h). We next quantified all Myosin light chain-2 (MLC-2v) positive nuclei in transversal sections that were isolectin B4 negative (to rule out false positives from the intermingled capillary network) (Fig. 4i). In line with their immature phenotype, *Acomys* CMs exhibited increased proliferation compared to *Mus* (3.6% ± 0.5 vs 1.2% ± 0.07) (Fig. 4i). Overall, we report an increased proliferative potential of the ventricular myocardium to respond to ischemia in *Acomys*, likely due to the presence of less differentiated cardiomyocytes. Together with increased vascularization of the scar and improved cardiac function after MI, these results demonstrate that *Acomys* presents a unique example of ischemic tolerance and cardiac repair in the mammalian class and provide a new experimental paradigm for cardiac research.

## Discussion

Given the crushing medical complications of patients with end-stage heart failure, the marked global increase in prevalence, and the severe limitation of donor hearts for transplantation, heart regeneration research has been the most compelling case for translating fundamental advances in stem cell science into clinical practice. Yet, as of today, an applicable working therapy to human patients still awaits discovery, urging the need for new and promising avenues in regeneration biology. The essence of regeneration biology comes from our current regime of cardiac models, including zebrafish, teleost fish, urodele amphibians, and neonatal mice and pigs ^19^. While these models have significantly advanced our understanding of regeneration biology, and in some cases have led to marked improvements in cardiomyocyte expansion in adult mice ^20–23^, these models impose restrictions in that they are either of non-mammalian vertebrate (e.g. zebrafish or newts), or neonatal vertebrate (mice or pigs) origin. On the basis of our findings in this study, we propose a new mammalian model for adult cardiac repair; the African spiny mouse. Compared to adult C57BL/6 mice, we demonstrate superior survivability in spiny mice, with accompanying near-absence of ventricular remodeling. In human patients, ventricular remodeling still occurs in around 30% of human myocardial infarcts despite timely primary coronary intervention and optimal standard pharmacotherapy (e.g. angiotensin-converting enzyme inhibitors (ACE-I), β-blockers, etc.) ^24^.

Interestingly, our findings revealed that scar size area correlates poorly with ejection fraction, contrary to *Mus*, which may suggest that *Acomys* scars hold properties that can foster regeneration rather than pathological remodeling. Notably, ventricular conductance only showed mild alterations, with no significant changes in Q wave dynamics. In infarcted hearts of human patients, pathological Q waves are characteristic of necrotic tissue and represent the area of the myocardium that cannot be depolarized. How *Acomys* hearts maintain conductivity in the presence of scar tissue, and whether or not the scar itself allows electrical conductance warrants further investigation, and may offer a new and currently unexplored platform for prevention strategies towards ventricular remodeling and heart failure.

In addition, we show partial functional recovery in *Acomys*, possibly owing to increased vascularization and cardiomyocyte expansion, with an associated improvement in heart function. Whether *Acomys* can completely regenerate the heart, including the resolution of existing scar tissue and full reconstitution of the myocardium remains to be determined, but may require prolonged periods of time that are reflective of the size and complexity of adult mammalian hearts. Nonetheless, its mammalian background presents unique opportunities for cardiac research that permits the study of a more systems-based approach within the context of mammalian physiology for the first time. Future studies will have to further characterize the regenerative repertoire of *Acomys* within the context of heart repair, ultimately working towards a druggable (set of) targets that can be exploited in the clinic to promote cardiac health.

## Acknowledgements

We thank the Utrecht Sequencing Facility for sequencing and Anko de Graaff and the Hubrecht Imaging Centre for assistance with microscopy. We also thank Dennis de Bakker and Sarah Maan Kamel for providing the zebrafish hearts. This work was supported by an ERC starting grant (granted to KB) (716894).

## Author contributions

KB and TK designed and oversaw the study. TK performed the bulk of the experiments and data analysis, prepared all figures and wrote the manuscript with comments from ER, KB, and EvR. HB processed the bulk sequencing samples for sequencing and performed alignment and counting. ER processed and performed the RT-qPCR experiments. DM quantified the MLC-2v immunostainings. AT performed the MLC-2v and Isolectin B4 immunostainings. JB wrote the ECG script. DV performed all heart surgeries. EvR provided expertise in experimental set-up and data interpretation.

## Methods

### Animals

All animal experiments were conducted under strict governmental and European guidelines and were approved by the Animal Welfare Committee of the Royal Netherlands Academy of Arts and Sciences, under license number AVD80100 2018 7144. Adult pathogen-free male C57BL/6 mice (7-9 weeks old) were obtained from Charles River Laboratories and group-housed at a density of 2-4 individuals in static individually ventilated cages (IVC) (Tecniplast, #GM500) with corncob granules (Rehofix, #MK1500). They were given autoclaved water and high nutrient food chow (RM3 pellets, Special diet) *ad libitum*. Adult pathogen-free male African spiny mice (7-9 weeks old) were bred and maintained in-house. They were group housed at a density of 2-8 individuals in static IVC (Tecniplast, #GR1800DD) with powdered cellulose pellets (ARBOCEL) and given autoclaved water and low nutrient and low protein food chow (RM1 pellets, Special diet) *ad libitum*. On occasion, and always after surgery, they additionally received black-oil sunflower seeds and a combination of dried vegetables. All cages were kept in climate-controlled quarters with a 12h/12h light/dark cycle.

### LAD ligation surgery

Myocardial infarction, ischemia-reperfusion, or sham surgery was performed in adult mice through permanent or temporary ligation (60 min) of the left anterior descending artery (LAD). Mice were anesthetized by an intraperitoneal injection of a Fentanyl (50 μg/kg), Midazolam (5 mg/kg), and Dexmedetomidine (500 μg/kg) cocktail, hereafter referred to as FMD. Monitoring anesthetic depth was assessed by toe reflex. Animals were placed on a warming plate (39 °C) and eyes were covered with Bepanthen to avoid dehydration. The thorax was shaved and disinfected with betadine and sterile phosphate buffered saline (PBS). A tracheal tube was placed and the mouse was connected to a ventilator (UNO Microventilator UMV-03, Uno BV.). Using aseptic technique with sterile instruments the skin was incised at the midline to allow access to the left third intercostal space. Pectoral muscles were retracted and the intercostal muscles cut caudal to the third rib. Wound hooks were placed to allow access to the heart. The pericardium was incised longitudinally and the LAD was identified. A 7.0 silk suture was placed around the LAD for permanent blocking of arterial blood flow. For the ischemia-reperfusion model, an additional piece of 2-3 mm PE 10 tubing was placed between the suture. After 60 min, the tubing was removed and the ligature cut to allow for reperfusion via the LAD. After surgery, mice were injected with 0.1 mg/kg of Buprenorphine for Mus and 200 mg/kg Metamizole for Acomys, after which the rib cage as well as skin were closed with a 5.0 silk suture. For C57BL/6 mice, the anesthetic cocktail was antagonized through an i.p. injection containing Atipamezol (2 mg/kg). For African spiny mice, anesthesia was antagonized through an i.p. injection containing Atipamezol (2 mg/kg), Flumazenil (0.1 mg/kg), Naloxone (0.6 mg/kg). The animal was disconnected from the ventilator by removing the tracheal tube and placed on a nose cone with 100% oxygen. When the animal was capable of autonomous breathing, it was moved to its cage (placed on a 40 °C heating mat) for further recovery. Subsequent pain relief during the following days was achieved through the addition of Metamizol (160 mg/kg) to the drinking water, stored in light-protected sterile bottles, for a maximum of 4 days.

### Echocardiography

Cardiac function was determined by two-dimensional transthoracic echocardiography on sedated mice (1.5-2.0% isoflurane), using a a 30 MHz transducer operated by a Vevo 2100 ultrasound scanner (FUJIFILM VisualSonics Inc.). Based on M-mode tracings, morphometric parameters, e.g. LVID;d, LVID;s, LVFW;s, LVFW;d, IVS;s and IVS;d, were measured according to the American Society of Echocardiography leading edge rule ^25^. These parameters were averaged based on five measurements. FS (%) and EF (%) were calculated from the measured parameters as: ((LVID;d-LVID;s)/LVID;d)×100%; and ((LVID;d3−LVID;s3)/LVID;d3)×100%, respectively, using Vevo LAB image analysis software.

### Electrocardiography (ECG)

Electrocardiographs (ECG) were obtained during echocardiography measurements by the Vevo 2100 Imaging System. For the analysis of QRS amplitude and duration, a custom script was written using .NET Core 3.1 (C# 8.0). QRS-complexes were automatically detected using peak detection software. R peak detection was based on a previously published algorithm to ensure accurate detection in severely affected infarcted hearts characterized by abnormal ECG signatures ^26^. Amplitudes were determined relative to the electromagnetic neutral point. Peak durations were determined by the difference in time between the start of the peak and the end of the peak. Datasets were manually verified for correct ECG measurement and accurate peak annotation.

### Tissue collection

Mice were euthanized through 5% isoflurane followed by cervical dislocation, and the chest was opened to expose the heart. After cutting the right atrium, the heart was immediately perfused slowly by injecting 5-10 mL cold perfusion buffer (130 mM NaCl, 5 mM KCl, 0.5 mM NaH2PO4, 10 mM HEPES, 10 mM glucose, 20 mM 2,3-butanedione monoxime (Sigma, #31550), 0.1 mM Blebbistatin (Sigma, # B0560), 10 mM taurine, and 5 mM EDTA, adjusted to pH 7.8) into the left ventricle. After perfusion, the heart was removed and washed in cold perfusion buffer before further processed.

### Imaging

#### 2D imaging of tissue sections

Upon organ excision, organs were fixed overnight at 4 °C in 2% formaldehyde. The next day, fixed tissues were washed three times in phosphate buffered saline, embedded and frozen in optimal cutting temperature compound (OCT) (Leica, #14020108926) and stored at −20 °C. Transversal cross-sections of 8 µm were cut using a Cryostar NX70 cryostat (Thermo Scientific). For the staining protocol, sections were permeabilized in 0.1% Triton-X-100 in PBS for approximately 30 min at RT, and then washed with PBS. Sections were then blocked for non-specific binding with 1% BSA and 10% donkey serum (Abcam, #ab7475) in PBS for 60 minutes at RT, and then incubated with primary antibody in 1% BSA and 5% donkey serum in PBS, O/N at 4 °C (see Table 1). The next day, following washing, sections were incubated in PBS with fluorescent secondary antibody, for 120 min at RT in the dark. Finally, sections were washed and incubated with Hoechst 33342 nucleic acid stain (Invitrogen, #H1399) or fluorescent dye (see Table 1) for 15 min, washed in PBS, mounted with ProLong™ Gold Antifade (Thermo Scientific, #P36934), and stored at RT in the dark until solidified. Samples were then imaged immediately or stored until further analysis at 4 °C in the dark. Samples were imaged using a Leica Thunder Imager.

**Table 1.** Antibody and fluorescent dye list

| Primary antibodies | Company | Catalog number | Working dilution |
| --- | --- | --- | --- |
| α-SMA-647 | Novus Biologicals | NBP2-34522AF647 | 1:100 |
| MLC-2v | Synaptic Systems | 310 003 | 1:200 |
| **Secondary antibodies** |  |  |  |
| Goat anti-Rabbit IgG (H+L)-488 | Invitrogen | A-11034 | 1:500 |
| **Fluorescent dyes** |  |  |  |
| Isolectin GS-IB4-568 | Invitrogen | I21412 | 1:50 |
| Wheat Germ Agglutinin-488 | Invitrogen | W11261 | 1:200 |
| Hoechst 33342 | Invitrogen | H3570 | 1:1000 |

#### 3D imaging of whole mount tissue samples

Upon organ excision, organs were fixed overnight at 4 °C in 2% formaldehyde. The next day, fixed tissues were washed three times in phosphate buffered saline, and stored at RT in PBS containing 0.2% gelatin (Sigma Aldrich, #G1393), 0.5% Triton X-100 (Sigma Aldrich, #X100) and 0.01% Thimerosal (Sigma Aldrich, #T8784) (PBS-GT) for at least 24 hours. Samples stored in PBS-GT were incubated with pre-conjugated primary antibody in PBS-GT while shaking, for 36 hours at RT (see Table 1). Excessive antibody was removed by thorough washing in PBS-GT for 24 hours and refreshing the solution every 3 hours during the day. Samples were then cleared for 36 hours at RT with Vectashield (Vector, #H-1000) after which they were imaged immediately or stored at 4 °C in the dark. Samples were imaged in 35 mm glass-bottom dishes (VWR, #75856742) using a laser scanning confocal microscope (Leica SP8) whilst submerged in Vectashield.

#### In vivo EdU labeling

Animals were injected intraperitoneally with 100 uL solution containing 1 mg/mL EdU (Invitrogen, #A10044) dissolved in sterile physiological saline, 5, 3, and 1 day(s) before the point of sacrifice. For the 4-day time point the 5 day injection point was skipped. Following organ excision and fixation overnight in 2% formaldehyde, EdU was visualized using the Click-iT™ EdU Alexa Fluor™ 647 imaging kit (Invitrogen, #C10340), according to the manufacturer’s instructions. The Click-iT® reaction cocktail was incubated with the samples for 2 hours at RT. Tissues were then further processed according to the imaging protocol (see ‘*2D imaging of murine tissue sections’*).

#### Masson’s Trichrome histological staining

To visualize deposited matrix, Masson’s trichrome staining was performed (Sigma Aldrich, #HT15). In brief, samples were incubated for 15 min in Bouin’s solution (Sigma Aldrich, #HT10132) at 56 °C (water bath), and then washed under running tap water for 1 min. Samples were then immersed in Weigert’s iron hematoxylin solution (Sigma Aldrich, #HT1079) for 5-10 min (depending on freshness), and again washed under running tap water for 2 min. Samples were incubated with Briebrich Scarlet-Acid Fuchsin solution for 5 min, rinsed in dH_2_O, and incubated with Phosphotungstic / Phosphomolybdic acid solution for 5 min. Finally, samples were immersed in Aniline Blue solution for 10 min, washed in 1% acetic acid for 2 min, and further washed with dH_2_O, and then dehydrated through an ethanol gradient. Samples were then dipped 15 times and cleared in Roti®-Histol (Carl Roth, #6640) and mounted with Roti®-Histokitt (Carl Roth, #6638).

### Cardiomyocyte isolation and nuclei number / cell surface determination

Ventricular tissues were fixed in 4% formaldehyde for 24 hours at 4 °C, followed by incubation in 50% w/v potassium hydroxide solution (Alfa Aesar, #35621) for 16 hours at 4 °C under constant agitation. Zebrafish ventricles were only incubated in potassium hydroxide for 3 hours at RT. Tissues were gently crushed with tweezers to release dissociated cardiomyocytes, and then pipetted up and down using a 5 mL stripette for approximately 1 minute. Cells were washed in PBS, spun down at 300 g for 5 min, liquid was aspirated, and cells were again washed with PBS for a total of three times. Cells were then incubated with Hoechst 33342 nucleic acid stain (Invitrogen, #H1399) in PBS for 15 minutes, washed one more time, and finally suspended and mounted on a glass slide with ProLong™ Gold Antifade (Thermo Scientific, #P36934). Cells were imaged using a Leica Thunder Imager and Leica Image Files (LIFs) were imported directly in Fiji (ImageJ 1.53c, USA). Nuclei numbers were counted manually, and corresponding surface area was determined by multiplying the width and length of every cell.

### Proteomic profiling

Whole heart ventricles were snap frozen in liquid nitrogen, and then grinded using a mortar and pestle. Approximately 20 mg of tissue per sample was used for further analysis. Tissues were incubated with 100 µL RIPA lysis buffer (65 mM Tris-HCl, 150 mM NaCl, 1% Triton-X-100, 1% Sodium deoxycholate, 1 mM EDTA, pH 7.4), and a mixture of protease inhibitors using cOmplete™ ULTRA Tablets (Roche, #5892791001). Lysates were kept on ice for 15 min, vortexed, and subsequently sonicated for 3 cycles (30 seconds ON, 30 seconds OFF) on low settings using a Bioruptor® Plus (Diagenode, #B01020001). Samples were then centrifuged for 10 min at 10.000 g and supernatant was collected. A BCA protein assay kit (Thermo Scientific, #23225) was used to determine protein content of the supernatant fractions and 300 µg of total protein was used for the following steps. Angiogenic proteins were measured using a Mouse Angiogenesis Array Kit (R&D Systems, #ARY015) according to the manufacturer’s instructions. Digital images were quantified by densitometry using LI-COR Image Studio Lite software (version 5.2.5).

### Quantitative real-time PCR

Whole heart ventricles were snap frozen in liquid nitrogen, and then grinded using a mortar and pestle. Total RNA was isolated from heart ventricles using a NucleoSpin RNA Plus isolation kit (Macherey-Nagel, #740984) according to the manufacturer’s instructions, and eluted in nuclease-free water (Invitrogen, #AM9937). Total RNA yield was determined with a NanoDrop ND-1000 spectrophotometer and samples were normalized accordingly. cDNA was synthesized using 300 ng total RNA per sample, using the RevertAid First Strand cDNA Synthesis Kit with random hexamer primers, according to the manufacturer’s instructions (Thermo Scientific, #K1622). The cDNA was diluted 30x with nuclease free water and used for quantitative PCR (qPCR). Each qPCR reaction was performed in a 10 uL volume, using 5 µL 2x iTaq universal SYBR® green supermix (Biorad, #1725124), 2 µL 2.5 µM target gene-specific forward and reverse oligo mixture and 3 µL cDNA. All primer sets were tested for linearity and single melting curves. A two-step qPCR program of 40 cycles was used with anneal/elongation step of 1 minute at 58 °C. All reactions were performed in technical duplicates and all targets were normalized to four housekeeping genes (18 S ribosomal RNA, Ribosomal Protein L13A (RPL13A), Ribosomal Protein L32 (RPL32), and Beta-2-Microglobulin (B2M) (*Acomys* only), or TATA-Box Binding Protein (Tbp) (*Mus* only). Analysis of RT-qPCR data was done using LinRegPCR analysis software ^27,28^. Gene units are expressed as the N0, which denotes the (unitless) RNA starting concentration. The N0 per sample is calculated in the unit of the Y-axis of the PCR amplification plot, which are arbitrary fluorescence units. Table 2 provides a list with all used primer sequences.

**Table 2.**
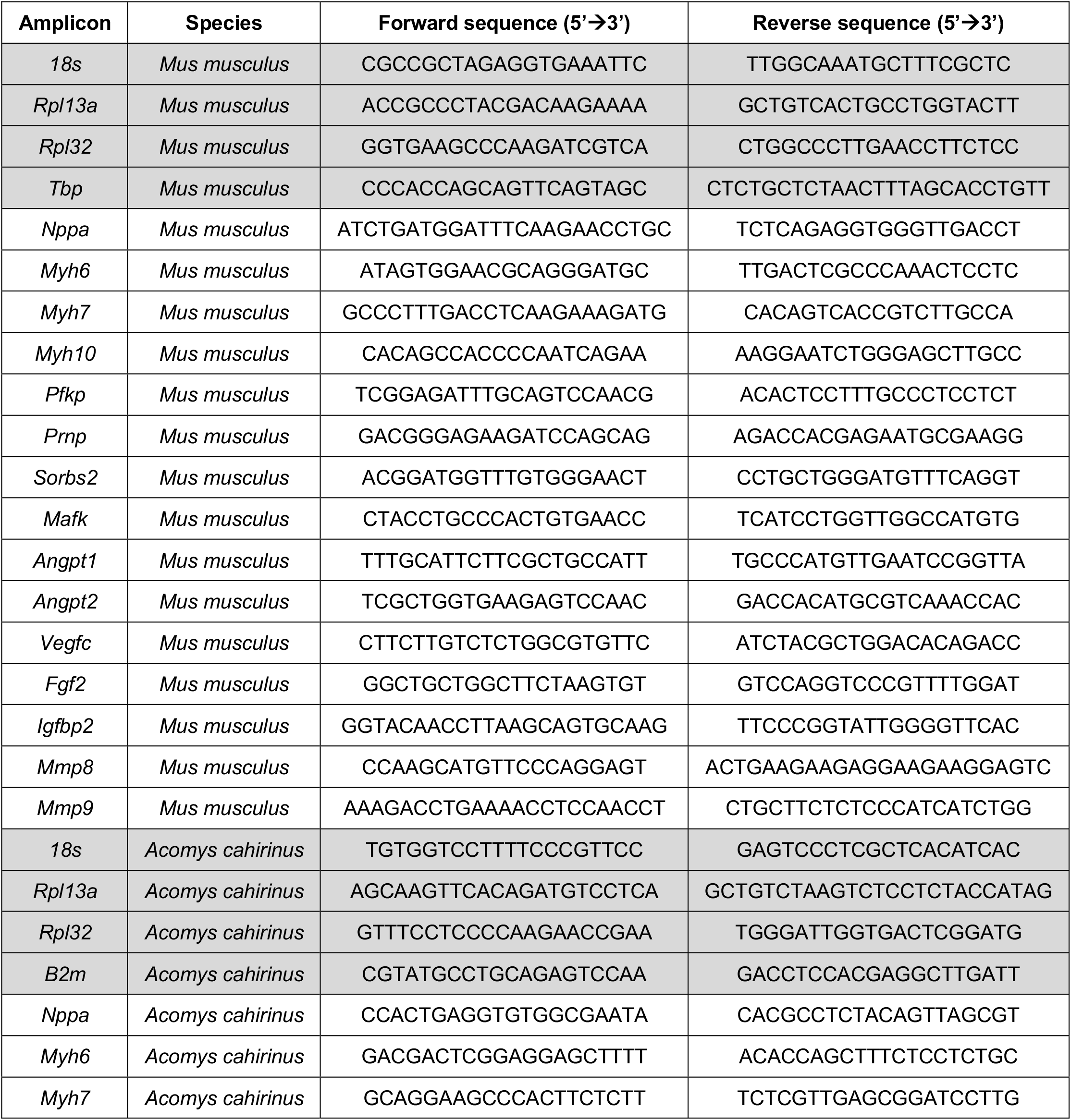
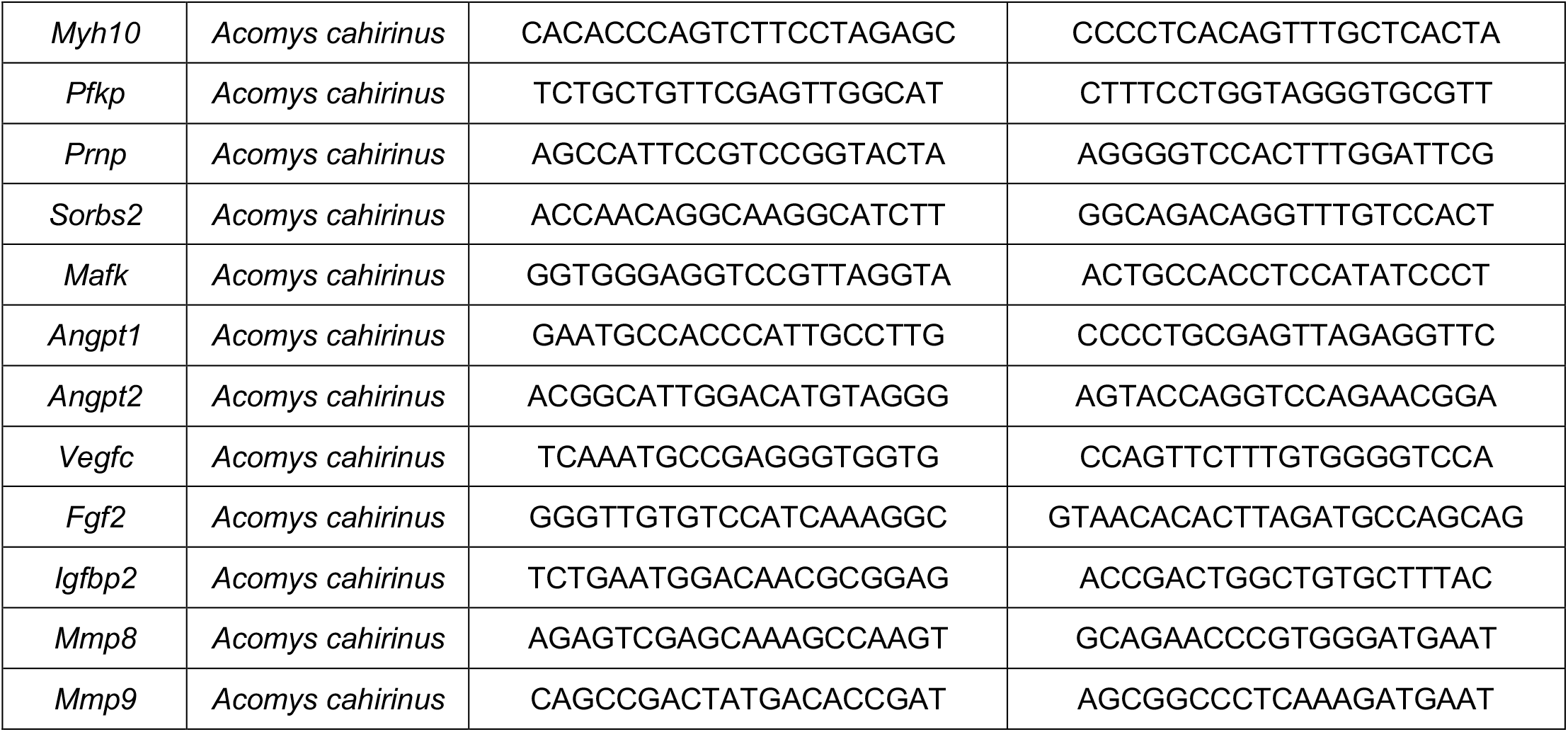
Primer sequences (housekeeping genes in grey)

### Image analysis

#### Scar area

Masson Trichrome histological stainings were used to mark the collagenous scar in injured hearts. Images were captured using a Leica DM4000 upright brightfield microscope, and source images were subsequently exported as TIFs. White balance was automatically corrected with the eye dropper function using Adobe Photoshop CC 2018, after which they were loaded in Fiji (ImageJ 1.53c, USA). To measure scar area, images were manually color thresholded based on Hue (140 to 200), Saturation (10 to 255), and Brightness (0 to 255) settings. The resulting threshold was selected and area was measured. To measure total heart area, the Hue settings were expanded (60 to 255) until the entire tissue section was thresholded. To account for changes in scar shape along the heart wall, six equally spaced-out transversal stacks, from point of ligation down to the tip of the apex, were used for image analysis (per animal), and the mean area was used to calculate scar area (relative to total area).

#### Heart size

For the analysis of heart size, the same set of images used to calculate scar area were used, and the corresponding mean area was used to calculate heart size. In brief, source images (Leica Imaging File) were loaded into Fiji (ImageJ 1.53c, USA), and were split into the different RGB channels. Using the green channel, images were Gaussian blurred (sigma = 50), and manually thresholded (“Triangle” algorithm) per batch (to account for differences in staining intensity between batches). The resulting threshold was converted to a mask (“Black Background” checked) and the particle was analyzed (“Exclude on edges”, and “Include holes” checked). A macro was written to automate the above steps for smooth processing.

#### General proliferation

In brief, source images (Leica Imaging File) were loaded into Fiji (ImageJ 1.53c, USA), and were split into the different fluorescent channels. Background was subtracted on all channels using the rolling ball algorithm (size = 50). To determine total nuclei and EdU+ nuclei numbers, the Hoechst or EdU channel was selected and a median filter (radius = 1) was applied. The image was subsequently thresholded using the Default (for Hoechst) or Yen (for EdU) algorithm. Particles were then analyzed (“Exclude on edges”, and “Include holes” checked) to count total nuclei numbers. To determine nuclei numbers inside the scar area, the WGA channel was selected, and the same macro was applied (Yen algorithm, but with a larger filter size (median filter, radius = 20). The resulting segmentation was added as a region of interest (ROI). EdU- and EdU+ nuclei were then counted inside the ROI.

#### Capillary proliferation

To determine the capillary area, the Isolectin B4 channel was selected, images were background subtracted using the rolling ball algorithm (size = 50), and then thresholded manually using the Moments algorithm to account for the higher Isolectin B4 expression in non-scar regions, therefore precluding automated thresholding. Strict criteria were used to avoid false positives. The resulting segmentation was added as a ROI, and the above settings for Hoechst, EdU, and WGA were applied to count the relevant proliferating capillaries in the scar area.

#### Cardiomyocyte proliferation

To detect proliferating cardiomyocytes (CMs), the MLC-2v channel was selected, background subtraction was performed using the rolling ball algorithm (size = 50) and a median filter was applied (size = 1) followed by automated thresholding (Default algorithm). To ensure there were no non-CM nuclei inside the MLC-2v segmentation, a subsequent mask containing all capillaries (which are normally closely intermingled with CMs) was generated that could then be subtracted from the MLC-2v segment. Because the scar region was not relevant here (due to absence of CMs), the Isolectin B4+ capillary area could be automatically thresholded (rolling ball algorithm of size 5 and median filter of radius 20) using the Default algorithm. These settings were less strict as the manual approach, to ensure a faithful exclusion of proliferating capillaries from the MLC-2v segment. Settings for Hoechst and EdU were then applied as described above. The Image calculator was finally used to determine nuclei numbers whose signal overlapped with that of Edu and CMs but not with Isolectin B4.

### Bulk sequencing

#### Pre-processing

Whole heart ventricles were snap frozen in liquid nitrogen, and then grinded using a mortar and pestle. Total RNA was isolated from approximately 20 mg of tissue using TRIzol Reagent (Invitrogen, #15596018). After RNA extraction, pellets were resuspended with barcoded primers (containing an anchored polyT, 6 bp unique barcode, 6 bp unique molecular identifier (UMI), 5’ Illumina adapter (used in the Illumina small RNA kit) and a T7 promoter). Barcode design was such that each pair differed by at least two nucleotides, so that a single sequencing error will not produce the wrong barcode. RNA samples were reverse transcribed to generate cDNA, pooled, and in vitro transcribed for linear amplification using the MessageAmp™ II aRNA Amplification Kit (Invitrogen, #AM1751) according to the CEL-seq protocol ^29^. Illumina sequencing libraries were prepared using the TruSeq Small RNA Library Prep Kit (Illumina, #RS-200), followed by PCR amplification for 14 rounds. Afterwards, libraries were sequenced paired-end at 26 bp read length for read 1 and 62 bp for read 2, using Illumina NextSeq500. Read 1 contained barcode information, whereas read 2 was used for alignment. The obtained reads were aligned to the mouse transcriptome (GRCm39) using Burrows-Wheeler Aligner’s Smith-Waterman alignment ^30^.

#### Differential expression analysis

After obtaining the raw read count expression matrix, samples were normalized and further processed in R using the DESeq2 Bioconductor package (version 3.12) ^31^. Values were transformed using regularized-logarithm transformation (Rld), and log2-transformed fold changes were shrunk using the apeglm method. An additive model was employed to compare the effects of age, while controlling for species. After differential expression analyses between relevant samples, the DESeq2 results table was exported for analysis using secondary platforms.

#### Gene set enrichment analysis

Differential analysis data from the DESeq2 pipeline was used for enrichment analysis, using the Bioconductor R package clusterProfiler ^32^. Gene symbols were mapped to EntrezIDs using the org.Mm.eg.db R package ^33^, and sorted in decreasing order based on the log(2) fold change. Genes were considered significantly differentially expressed if they had FDR-adjusted values of P < 0.05 and a log(2) fold chance threshold of 2.0 unless otherwise stated. The Reactome mouse database was used as a source of pathway data, using the Bioconductor ReactomePA R package ^34^. The *compareCluster* function was used to calculate and compare gene clusters between samples (up or down regulated enrichment terms).

#### Pairwise correlation matrix

Using R, we performed PCA on variable genes (variance > 0.5) expressed (>1 FPKM) in more than two samples. We extracted the genes correlating and anti-correlating with PC1-5, using an absolute PC loading threshold >0.2 with a maximum of 50 genes per principal component to avoid individual principal components swamping the analysis, resulting in 300 genes. To construct the network, we computed a pairwise correlation matrix for all samples, using genes discovered in the PCA analysis. We then generated a weighted adjacency network graph using the graph.adjacency() command in igraph and visualized samples as vertices connected to other samples via edges if the Pearson pairwise correlation between two cells was higher than 0.6. The Fruchterman–Reingold layout was used to plot the network graph.

#### Cell cycle prediction

Cell cycle prediction was applied to classify samples into G1, G2/M, or S phase ^18^ based on expression patterns of pre-trained classifiers. Results were generated as part of the single-cell R-analysis tools (SCRAT) pipeline ^35^.

### Statistical analyses

All data represent the mean ± standard error of the mean (SEM). Shapiro-Wilk and D’Agostino & Pearson test (p>0.05) were used to test whether samples were normally distributed (approximately), using GraphPad Prism version 7. Two group comparisons were made using an unpaired Student’s t-test for normally distributed data. Comparisons between three or more groups were performed using a one-way ANOVA followed by Tukey’s post hoc test for normally distributed data. A value of p < 0.05 was considered statistically significant, where * is p < 0.05, ** is p < 0.01, and *** is p < 0.001. Analyses were performed with GraphPad Prism version 7 (GraphPad Software, Inc.).

---

**Extended figure 1.**
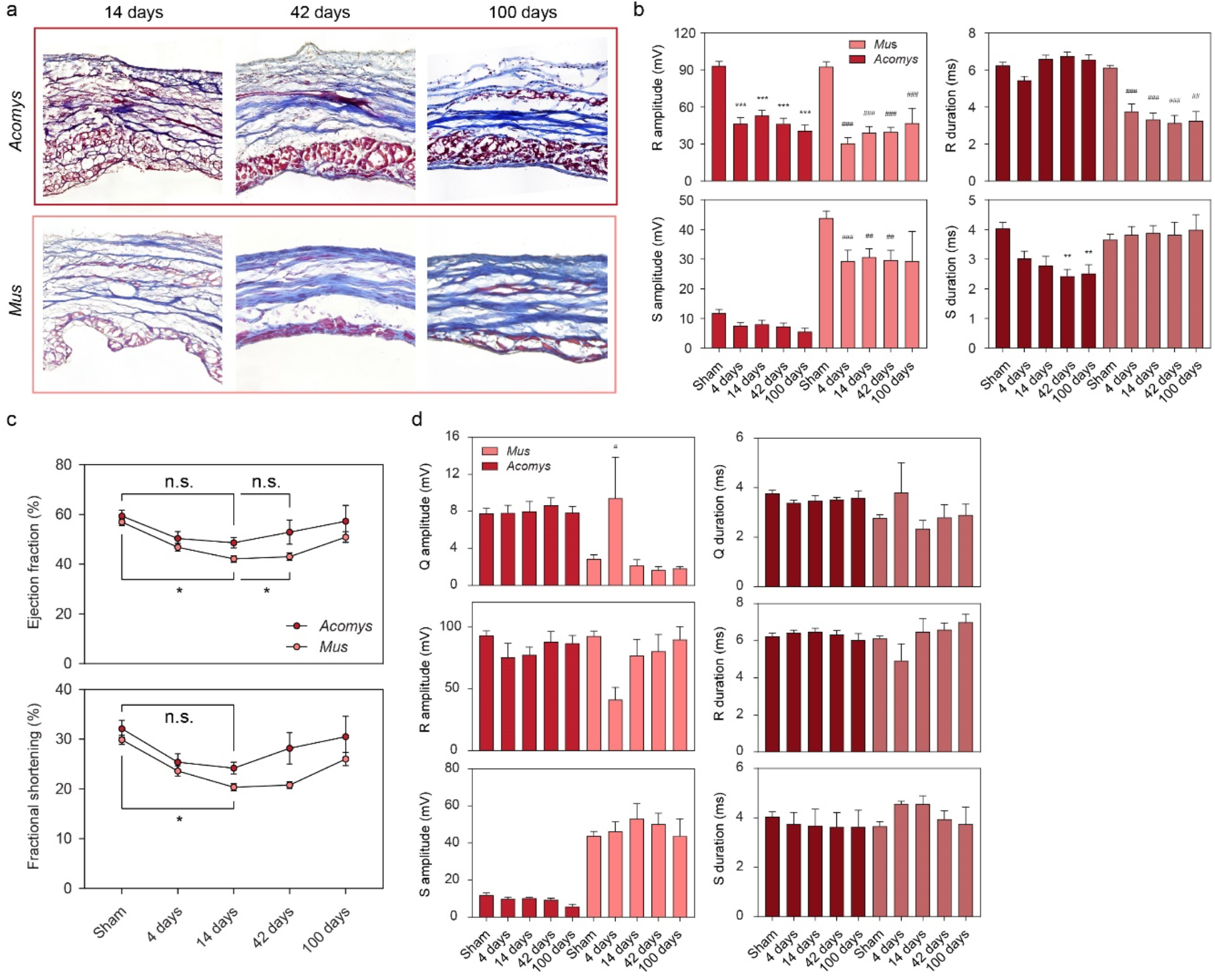
Ischemic tolerance in *Acomys* is substantiated by the reperfusion model. **(a)** Representative images of a Masson’s Trichrome stain showing the collagenous scar area after different time point. **(b)** Electrocardiogram quantification (see ‘Methods’) showing R and S wave parameters (amplitude and duration) after MI or Sham-control. * is compared to *Acomys*-Sham, # is compared to *Mus*-Sham. One-way ANOVA followed by Tukey’s post hoc test. n = 15 for *Acomys*, n = 14 for *Mus*. **(c)** Echocardiographic quantification of cardiac output after temporary ligation of the LAD for 60 min (reperfusion injury) or Sham-control, including ejection fraction (volumetric percentage of fluid ejected from the chamber with each contraction), and fractional shortening (percentage of diastolic dimension that is lost in systole). **(d)** Electrocardiogram quantification (see ‘Methods’) showing Q, R and S wave parameters (amplitude and duration) after reperfusion injury or Sham-control. # is compared to *Mus*-Sham. One-way ANOVA followed by Tukey’s post hoc test. * is p < 0.05, ** is p < 0.01, and *** is p < 0.001.

**Extended figure 2.**
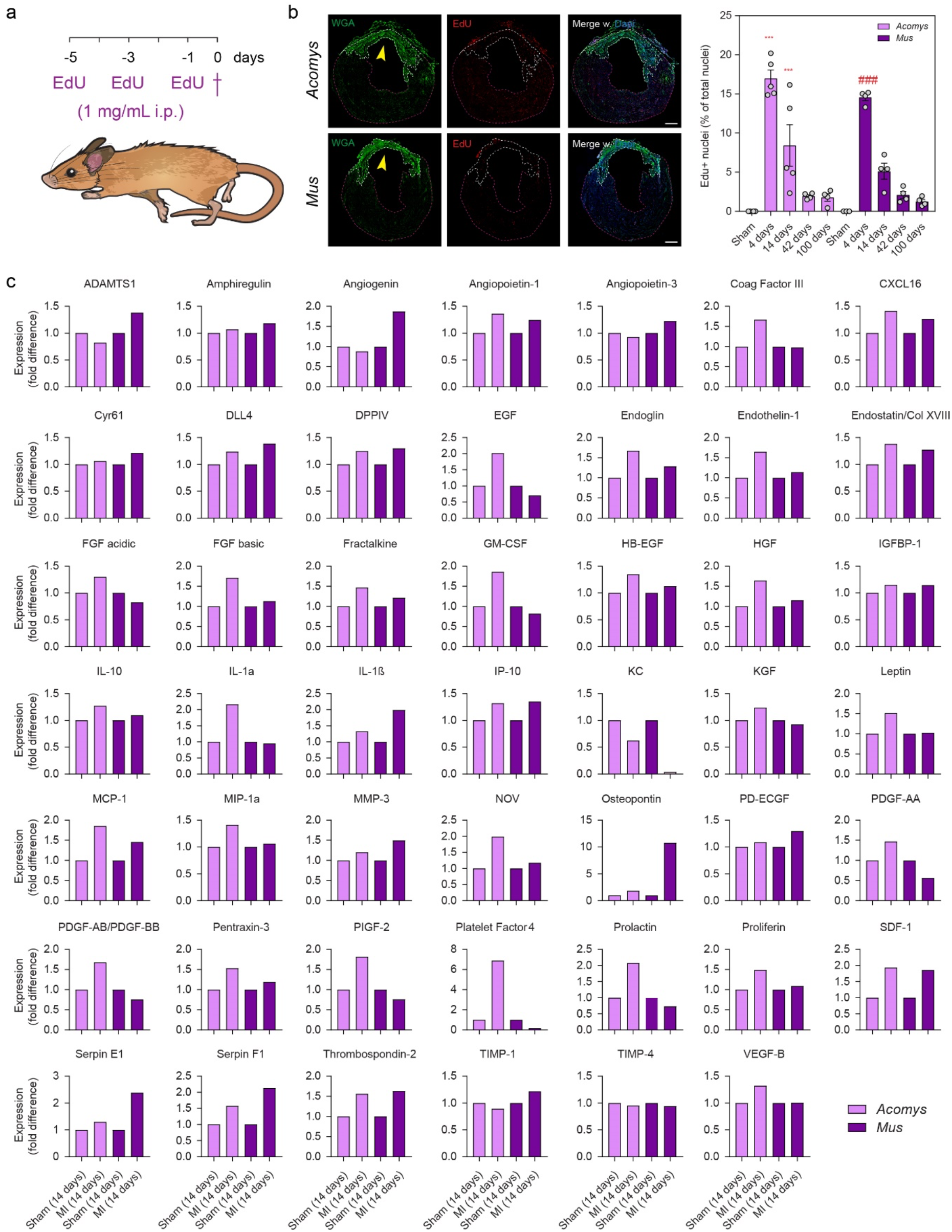
Expression of a large array of angiogenic proteins in injured *Acomys* hearts. **(a)** Overview of the EdU injection strategy. **(b)** Representative fluorescent image showing proliferation (marked by EdU labeling) in the scarred heart (yellow arrows) 14 days after MI, with corresponding quantification for all the time points. * is compared to *Acomys*-Sham, # is compared to *Mus*-Sham. One-way ANOVA followed by Tukey’s post hoc test. Scale bar = 700 µm. **(c)** Quantification from proteomic profiling of angiogenic proteins, obtained from whole heart ventricle homogenates (14 day post-MI and 14 day Sham-controls). Samples are normalized to the Sham-control (*Acomys* or *Mus*). * is p < 0.05, ** is p < 0.01, and *** is p < 0.001.

## References

1. Bui, A. L., Horwich, T. B. & Fonarow, G. C. Epidemiology and risk profile of heart failure. Nat. Rev. Cardiol. 8, 30–41 (2011).

2. Riehle, C. & Bauersachs, J. Small animal models of heart failure. Cardiovasc. Res. 115, 1838– 1849 (2019).

3. Kikuchi, K. et al. Primary contribution to zebrafish heart regeneration by gata4(+) cardiomyocytes. Nature 464, 601–605 (2010).

4. Jopling, C. et al. Zebrafish heart regeneration occurs by cardiomyocyte dedifferentiation and proliferation. Nature 464, 606–609 (2010).

5. Porrello, E. R. et al. Transient regenerative potential of the neonatal mouse heart. Science 331, 1078–1080 (2011).

6. Seifert, A. W. et al. Skin shedding and tissue regeneration in African spiny mice (Acomys). Nature 489, 561–565 (2012).

7. Gawriluk, T. R. et al. Comparative analysis of ear-hole closure identifies epimorphic regeneration as a discrete trait in mammals. Nat. Commun. 7, 11164 (2016).

8. Gaire, J. et al. Spiny mouse (Acomys): an emerging research organism for regenerative medicine with applications beyond the skin. Npj Regen. Med. 6, 1–6 (2021).

9. Shindo, S. et al. Evaluating spiny mice (Acomys) as a model for cardiac research. bioRxiv doi:10.1101/2020.09.29.317388.

10. Thygesen, K. et al. Universal definition of myocardial infarction. Circulation 116, 2634–2653 (2007).

11. Cohn, J. N., Ferrari, R. & Sharpe, N. Cardiac remodeling—concepts and clinical implications: a consensus paper from an international forum on cardiac remodeling. J. Am. Coll. Cardiol. 35, 569–582 (2000).

12. Vigil-Garcia, M. et al. Gene expression profiling of hypertrophic cardiomyocytes identifies new players in pathological remodeling. Cardiovasc. Res. (2020) doi:10.1093/cvr/cvaa233.

13. Felmeden, D. C., Blann, A. D. & Lip, G. Y. H. Angiogenesis: basic pathophysiology and implications for disease. Eur. Heart J. 24, 586–603 (2003).

14. Hirose, K. et al. Evidence for hormonal control of heart regenerative capacity during endothermy acquisition. Science 364, 184–188 (2019).

15. Patterson, M. et al. Frequency of mononuclear diploid cardiomyocytes underlies natural variation in heart regeneration. Nat. Genet. 49, 1346–1353 (2017).

16. Camp, J. G. et al. Human cerebral organoids recapitulate gene expression programs of fetal neocortex development. Proc. Natl. Acad. Sci. 112, 15672–15677 (2015).

17. Guo Yuxuan & Pu William T. Cardiomyocyte Maturation. Circ. Res. 126, 1086–1106 (2020).

18. Scialdone, A. et al. Computational assignment of cell-cycle stage from single-cell transcriptome data. Methods 85, 54–61 (2015).

19. Goldman, J. A. & Poss, K. D. Gene regulatory programmes of tissue regeneration. Nat. Rev. Genet. 21, 511–525 (2020).

20. Wei, K. et al. Epicardial FSTL1 reconstitution regenerates the adult mammalian heart. Nature 525, 479–485 (2015).

21. Nakada, Y. et al. Hypoxia induces heart regeneration in adult mice. Nature 541, 222–227 (2017).

22. Bassat, E. et al. The extracellular matrix protein agrin promotes heart regeneration in mice. Nature 547, 179–184 (2017).

23. Aharonov, A. et al. ERBB2 drives YAP activation and EMT-like processes during cardiac regeneration. Nat. Cell Biol. 22, 1346–1356 (2020).

24. Masci, P. G. et al. Relationship between location and size of myocardial infarction and their reciprocal influences on post-infarction left ventricular remodelling. Eur. Heart J. 32, 1640–1648 (2011).

25. Sahn D J, DeMaria A, Kisslo J & Weyman A. Recommendations regarding quantitation in M-mode echocardiography: results of a survey of echocardiographic measurements. Circulation 58, 1072–1083 (1978).

26. Sadhukhan, D. & Mitra, M. R-Peak Detection Algorithm for Ecg using Double Difference And RR Interval Processing. Procedia Technol. 4, 873–877 (2012).

27. Ruijter, J. M. et al. Amplification efficiency: linking baseline and bias in the analysis of quantitative PCR data. Nucleic Acids Res. 37, e45 (2009).

28. Ruijter, J. M. et al. Evaluation of qPCR curve analysis methods for reliable biomarker discovery: bias, resolution, precision, and implications. Methods San Diego Calif 59, 32–46 (2013).

29. Hashimshony, T., Wagner, F., Sher, N. & Yanai, I. CEL-Seq: Single-Cell RNA-Seq by Multiplexed Linear Amplification. Cell Rep. 2, 666–673 (2012).

30. Li, H. & Durbin, R. Fast and accurate long-read alignment with Burrows–Wheeler transform. Bioinformatics 26, 589–595 (2010).

31. Love, M. I., Huber, W. & Anders, S. Moderated estimation of fold change and dispersion for RNA-seq data with DESeq2. Genome Biol. 15, 550 (2014).

32. Yu, G., Wang, L.-G., Han, Y. & He, Q.-Y. clusterProfiler: an R Package for Comparing Biological Themes Among Gene Clusters. OMICS J. Integr. Biol. 16, 284–287 (2012).

33. Carlson, M. org.Mm.eg.db: Genome wide annotation for Mouse. (2019).

34. Yu, G. & He, Q.-Y. ReactomePA: an R/Bioconductor package for reactome pathway analysis and visualization. Mol. Biosyst. 12, 477–479 (2016).

35. Camp, J. G. et al. Multilineage communication regulates human liver bud development from pluripotency. Nature 546, 533–538 (2017).

